# The riboflavin biosynthetic pathway as a novel target for antifungal drugs against *Candida* species

**DOI:** 10.1101/2024.03.01.582991

**Authors:** Jana Nysten, Arne Peetermans, Dries Vaneynde, Liesbeth Demuyser, Patrick Van Dijck

**Author notes:** Address correspondance to Patrick Van Dijck,. Department of Biology and Biological Engineering, Industrial Biotechnology, Chalmers University of Technology, Gothenburg, Sweden (Arne Peetermans). Department of Microbial and Molecular Systems, KU Leuven, Leuven, Belgium.

## Abstract

In recent decades, there has been an increase in the occurrence of fungal infections, yet the arsenal of drugs available to fight invasive infections remains very limited. The development of new antifungal agents is hindered by the restricted number of molecular targets that can be exploited, given the shared eukaryotic nature of fungi and their hosts which often leads to host toxicity. In this paper, we examine the riboflavin biosynthetic pathway as a potential novel drug target.

Riboflavin is an essential nutrient for all living organisms. Its biosynthetic pathway does not exist in humans, who obtain riboflavin through their diet. Our findings demonstrate that all enzymes in the pathway are essential for *Candida albicans*, *Candida glabrata,* and *Saccharomyces cerevisiae.* Among these enzymes, Rib1 and Rib3 are the most promising targets. Auxotrophic strains, which mimic a drug targeting the biosynthesis pathway, experience rapid mortality in the absence of supplemented riboflavin. Nevertheless, the cells can still take up external riboflavin when supplemented. We identified Orf19.4337 as the riboflavin importer in *C. albicans* and named it Rut1. We found that Rut1 only facilitates growth at external riboflavin concentrations that exceed the physiological concentrations in the human body, making it unlikely that riboflavin uptake to act as a potential resistance mechanism for a drug targeting the biosynthesis pathway. Interestingly, the uptake system in *S. cerevisiae* is more effective than in *C. albicans* and *C. glabrata,* enabling an auxotrophic *S. cerevisiae* strain to outcompete an auxotrophic *C. albicans* strain in lower riboflavin concentrations.

**Importance:** *Candida* species are a common cause of invasive fungal infections. *Candida albicans,* in particular, poses a significant threat to immunocompromised individuals. This opportunistic pathogen typically lives as a commensal on mucosal surfaces of healthy individuals, but it can also cause invasive infections associated with high morbidity and mortality. Currently, there are only three major classes of antifungal drugs available to treat these infections. Additionally, the efficacy of these antifungal agents is restricted by host toxicity, suboptimal pharmacokinetics, a narrow spectrum of activity, intrinsic resistance of fungal species, such as *Candida glabrata*, to certain drugs, and the acquisition of resistance over time. Therefore, it is crucial to identify new antifungal drug targets with novel modes of action to add to the limited armamentarium.

## Introduction

Fungal infections affect around 1.7 billion people annually, making it a global health concern (1). Alarmingly, this number is continuously rising due to the growing population of at-risk patients, ironically arising from advances in modern medicine aiming to expand the human lifespan (2). According to recent estimates, fungal infections cause over 3.752 million overall deaths and 2.548 attributable deaths each year with *Candida*, *Aspergillus*, *Pneumocystis,* and *Cryptococcus* genera being the primary causative agents (1, 3, 4). *Candida albicans* is a leading cause of nosocomial infections and has a mortality rate of about 63.6%, prompting its recent classification by the World Health Organization as a pathogen that is of critical concern to public health and requires immediate attention (4–6). This opportunistic pathogen colonizes the gastrointestinal tract, skin, mouth, and genitourinary tract of over 70% of the world’s population (7). Although this pathogen typically maintains a commensal and asymptomatic relationship with its human host, it can become pathogenic under favorable conditions, resulting in clinical manifestations ranging from superficial mucosal infections to life-threatening invasive candidiasis (8). In addition to *C. albicans*, non-*albicans Candida* species like *Candida glabrata* (recently renamed as *Nakaseomyces glabratus)* and *Candida auris* are becoming more prevalent. These species have inherent resistance or possess a heightened ability to acquire resistance to commonly used antifungal drugs (9–11). This issue is compounded by the limited number of antifungal drugs available, with only three major classes of agents for treating invasive mycoses. Developing new antifungal drugs is challenging due to the shared eukaryotic nature of fungal and human host cells, resulting in a scarcity of unique molecular targets amenable to drug development (12, 13). The development of antimycotic drugs often encounters hurdles such as host toxicity, fungistatic activity, and the escalating crisis of antifungal drug resistance (14). Therefore, there is a pressing need for new drug targets with novel and distinct modes of action (15).

In this manuscript, we explore the riboflavin biosynthetic pathway as a potential antifungal drug target. Riboflavin, also known as vitamin B_2_ is a water-soluble micronutrient crucial for the vitality of all living organisms (16). Although many bacteria, fungi, and plants can synthesize this vitamin themselves, mammals, including humans, have lost this ability and rely on external sources such as nutrition and, to a lesser extent, intestinal microorganisms (17, 18). The synthesis of riboflavin in bacteria, fungi, and plants involves the conversion of one molecule of guanosine-5’-triphosphate (GTP) and two molecules of ribulose 5-phosphate by various Rib enzymes, ultimately forming riboflavin (19). The riboflavin biosynthetic pathway is depicted in Supplementary Figure 1. Riboflavin is subsequently converted into flavin mononucleotide (FMN) and flavin adenine dinucleotide (FAD). These two essential redox cofactors function as prosthetic groups for flavoproteins which catalyze redox reactions in nearly all metabolic pathways in prokaryotic and eukaryotic cells (20, 21). They play important roles in various cellular processes including β-oxidation, glucose metabolism, oxidative stress defence, protein folding, DNA and RNA metabolism, gluconeogenesis, erythropoiesis, and corticosteroid generation (22–26). We consider enzymes in this pathway promising drug targets due to the anticipated minimal host toxicity, as humans lack the requisite or analogous enzymes for *de novo* synthesis (27).

We examined the pathway in *C. albicans* and *C. glabrata*, and also compared it to the non-pathogenic model organism *S. cerevisiae*. We found all Rib enzymes to be essential for growth in all studied organisms, which contradicts previous studies identifying Rib1 and Rib4 as non-essential in *S. cerevisiae* (28). Our observations identify *Ca*Rib1 and *Ca*Rib3 as the most attractive drug targets in the pathway. Additionally, riboflavin auxotrophic *C. albicans* and *C. glabrata* cells, mimicking a drug targeting the riboflavin biosynthetic pathway, experience rapid mortality within 24 hours in RPMI. In auxotrophic *C. albicans* cells, some growth over 72 hours can be achieved when 20 mg/L of riboflavin is added to the medium, with higher concentrations resulting in better growth. The significance of riboflavin biosynthesis as a promising antifungal drug target is underscored by the avirulence of a *Carib1Δ/Δ* strain in a systemic mouse model, as these cells cannot survive in physiological riboflavin concentrations (29). Additionally, we explore riboflavin import as a potential mechanism to counteract the mode of action of such a drug. Our study on *C. albicans* identifies Orf19.4337 as the primary riboflavin importer. We named it Rut1 for Riboflavin Uptake Transporter. However, even in the presence of high riboflavin concentrations of 200 mg/L, growth in auxotrophic strains cannot be fully restored to wild-type levels. This demonstrates a partial inefficiency of the transport system, leading us to speculate that import won’t be able to nullify a drug targeting the pathway.

Expanding our investigation to *S. cerevisiae*, we found the uptake system to be more effective than in the two *Candida* species, enabling better growth on low levels of riboflavin. Co-culturing auxotrophic *S. cerevisiae* and *C. albicans RIB1* deletion strains in media with 5 mg/L of riboflavin indicates that *S. cerevisiae* outcompetes *C. albicans*, whereas, under wild-type prototrophic conditions, *S. cerevisiae* grows more slowly. Our study suggests that drugs targeting the riboflavin biosynthetic pathway could be effective in inhibiting *C. albicans* and *C. glabrata* while potentially sparing *S. cerevisiae*. This has potentially interesting implications as *S. cerevisiae* has probiotic properties that can inhibit *C. albicans* in certain niches (30).

## Results

### All enzymes involved in riboflavin biosynthesis are essential for growth in *C. albicans*

To determine the essentiality of the various Rib enzymes in the riboflavin biosynthetic pathway, we generated homozygous deletion strains using CRISPR-Cas9. Figures 1A and 1B illustrate the growth profiles of the *RIB* deletion strains and the wild-type strain in regular LoFlo medium without supplementation of external riboflavin and in LoFlo medium where 200 mg/L of riboflavin was supplemented, respectively. It was observed that none of the *RIB* deletion strains were able to grow, except when external riboflavin was added to the medium. At concentrations approaching the maximum solubility level of 200 mg/L, growth was partially restored but not to wild-type levels, suggesting the presence of a riboflavin uptake system with limited effectiveness. To functionally validate our findings, we reintroduced the *RIB* genes into the corresponding deletion strains and assessed their performance in LoFlo media without riboflavin supplementation, as illustrated in Supplementary Figure 2A. The reintegrated genes successfully rescued the phenotype, aligning with our expectations. The phenotypes observed in LoFlo medium containing 2% glycerol as a carbon source instead of glucose and in YPD medium are similar to those obtained in LoFlo medium with glucose (Supplementary Figure 2B, C, D, E, F). As expected, all strains grew slower in glycerol than in glucose-containing medium.

**Figure 1:**
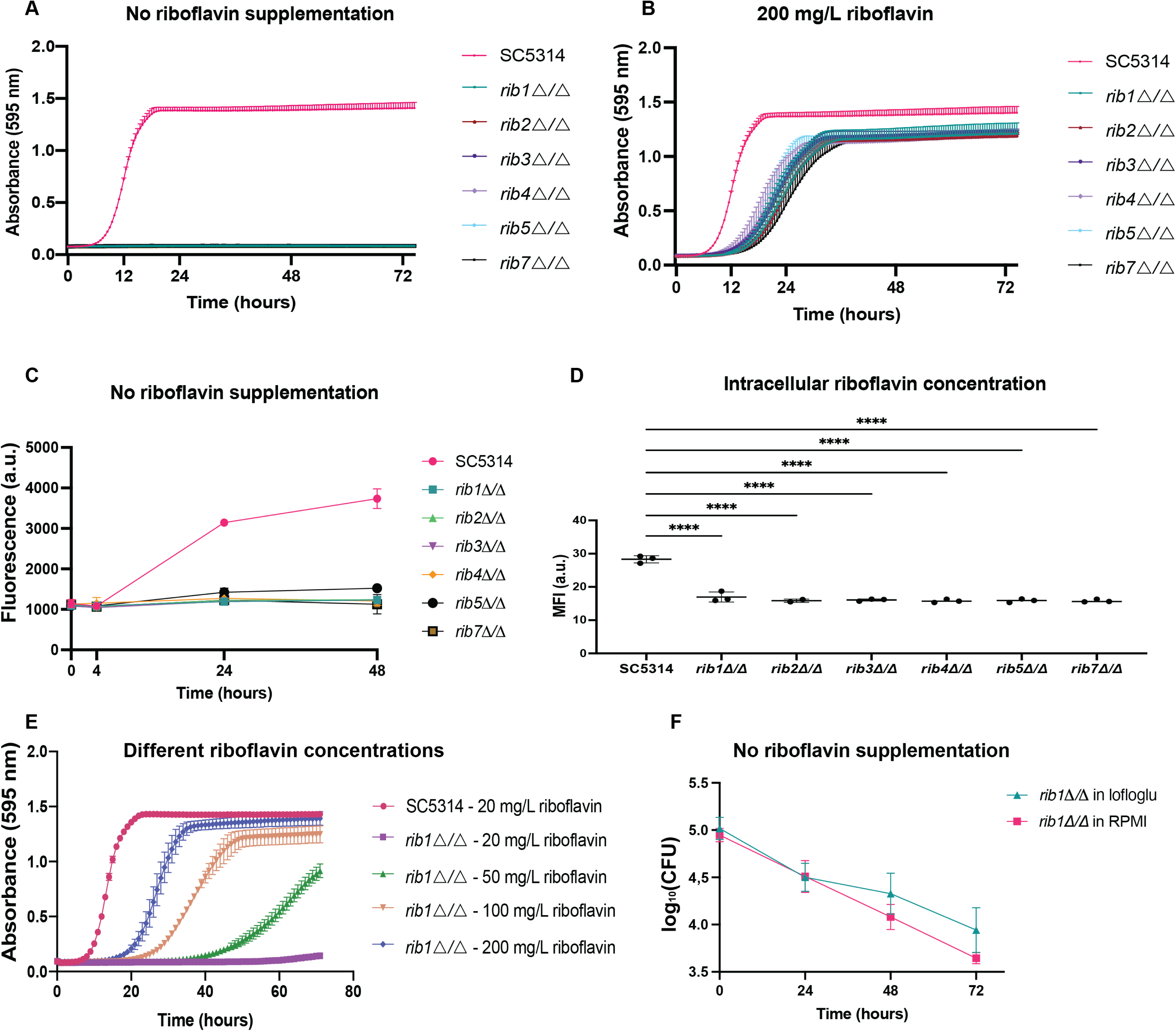
The crucial role of riboflavin biosynthetic enzymes in *C. albicans.* (A) Homozygous deletion strains of all enzymes in the riboflavin biosynthetic pathway were cultured in a growth medium without supplemented riboflavin, and their absorbance was measured over time. (B) Growth of strains in a medium supplemented with 200 mg/L of riboflavin. (C) Growth of strains in a growth medium without external riboflavin supplementation and riboflavin production measured by fluorescent measurements. (D) Intracellular riboflavin levels were measured by fluorescent measurements in a flow cytometer where 10,000 cells were monitored per biological repeat. (E) Growth of the *RIB1* deletion strain was monitored in LoFlo medium with varying concentrations of riboflavin supplementation. (F) Growth of a *RIB1* deletion strain in the absence of supplemented riboflavin in both LoFlo and in blood-simulative RPMI medium that contains a physiological riboflavin concentration of 0.2 mg/L. To assess cell survival, colony-forming units (CFUs) were counted every 24 hours on YPD medium with 200 mg/L riboflavin to facilitate the growth of the surviving auxotrophic strains. All experiments were conducted with three biological replicates, and the mean ± standard deviations (s.d.) are presented. Statistical analysis was performed by ordinary one-way ANOVA with Dunnett’s multiple comparisons test ****, P < 0.0001.

As expected, a reduction in riboflavin production was detected in the cell culture of *RIB* deletion strains (Figure 1C). Figure 1D depicts the intracellular riboflavin levels measured by flow cytometry and shows that the auxotrophic strains exhibit low intracellular riboflavin levels compared to the wild-type strain. Subsequently, we assessed the growth of the *RIB1* deletion strain under varying extracellular riboflavin concentrations and found that the addition of 20 mg/L of riboflavin resulted in a minor growth increase over 72 hours. Importantly, increasing external riboflavin concentrations were found to be correlated with increased growth (Figure 1E). Additionally, rapid cell death was observed in the auxotrophic cells when no external riboflavin was present as well as in blood-simulative RPMI medium with physiological riboflavin concentrations of 0.2 mg/L (see Figure 1F). These physiological concentrations of riboflavin do not appear to affect *C. albicans* survival. These findings emphasize the crucial role of riboflavin in maintaining cellular viability and the potential vulnerability of fungal cells with impaired riboflavin biosynthesis, supporting our pursuit of identifying the riboflavin biosynthetic pathway as a valuable target for antifungal drugs.

### Rib1 and Rib3 are attractive drug targets

To identify the most promising drug targets within the pathway, we used the *SAT1* flipper approach to generate heterozygous deletion strains (31). We hypothesized that the deletion of one allele of a rate-limiting enzyme would manifest as reduced fluorescence or growth. Intriguingly, all heterozygous strains exhibited the same growth rate as the wild-type strain on LoFlo medium without externally added riboflavin (Figure 2A). However, the *RIB1/rib1*Δ and *RIB3/rib3*Δ strains demonstrated a significantly reduced riboflavin production compared to the other strains (Figure 2B). This observation was also confirmed via flow cytometry where the intracellular riboflavin content was measured (Figure 2C). This indicates that Rib1 and Rib3 are the most interesting drug targets of the pathway. The positional significance of Rib1 and Rib3 further reinforces their critical roles in riboflavin synthesis as these are the first enzymes active in their respective branches of the pathway. This result does not only emphasize the pivotal roles of Rib1 and Rib3 and thus their amenability to serve as potential antifungal drug targets, but it also highlights the intriguing finding that *C. albicans* can exhibit comparable growth on LoFlo medium even when producing less riboflavin, suggesting an excess production of this essential molecule.

**Figure 2:**
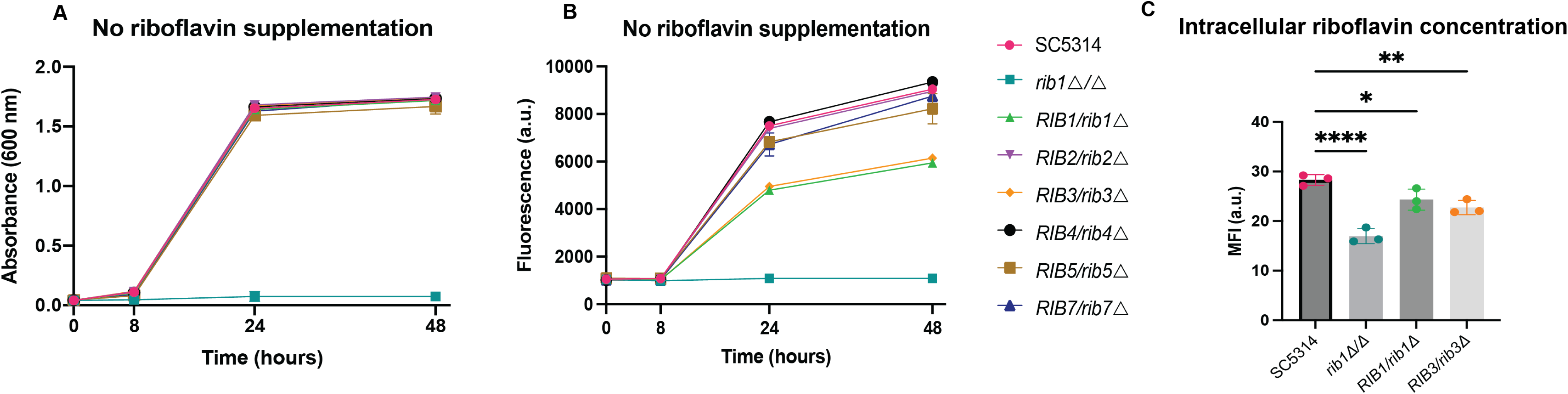
Rib1 and Rib3 are particularly attractive drug targets in the riboflavin biosynthetic pathway. (A) The cultivation of heterozygous *RIB* strains on LoFlo medium. Growth was monitored by measuring the absorbance at 600 nm. (B) The riboflavin content was measured spectrophotometrically. (C) Intracellular riboflavin levels were measured by fluorescent measurements in a flow cytometer where 10,000 cells were monitored per biological repeat. Statistical analysis was performed by ordinary one-way ANOVA with Dunnett’s multiple comparisons test *, P < 0.05; **, P < 0.01; ****, P < 0.0001. Each experiment was performed with three biological repeats and the mean ± s.d. is displayed.

### Rut1 (Orf19.4337) is a major riboflavin transporter in the human fungal pathogen *C. albicans*

Due to the absence of previous knowledge regarding riboflavin transporters in *C. albicans*, we endeavored to elucidate the transport mechanisms within this organism. We conducted this investigation by examining five potential riboflavin transporters, selected based on their orthology with *Sc*Mch5, the riboflavin importer in *S. cerevisiae* (28). To identify the true riboflavin importer in *C. albicans*, we deleted the four most prominent *ScMCH5* orthologous genes, based on protein-BLAST, in an auxotrophic *rib1Δ/Δ* background: *ORF19.6263*, *ORF19.5720*, *ORF19.1584*, *ORF19.2751*. However, none of these orthologs appear to have a role in riboflavin uptake as the deletion of these individual genes in the auxotrophic strain did not result in reduced growth as shown in supplementary Figure 3A. We hypothesized that functional redundancy among these transporters may underlie riboflavin uptake, wherein the deletion of one transporter could be compensated for by the functional takeover of another. Therefore, we deleted the four *ScMCH5* orthologous genes in the auxotrophic *rib1Δ/Δ* deletion background to generate a quintuple deletion strain. However, also here, we did not observe a change in phenotype. Gaur *et al*. proposed another potential riboflavin transporter, Orf19.4337 through computational analysis (32). This protein did not appear as a top hit in the initial pBLAST search as it has more sequence similarity with *Sc*Mch3 (*ScESBP6*) than with *Sc*Mch5. Subsequent experiments confirmed Orf19.4337 (C5_03080C_A) as the true riboflavin transporter, evidenced by the inability of the *rib1Δ/Δ* deletion strain to take up 50 mg/L or 100 mg/L external riboflavin in the absence of this gene (Figure 3A and 3B). Therefore, we have named this transporter Rut1, which stands for Riboflavin Uptake Transporter, and we will refer to this gene as Rut1 throughout the rest of this paper. Furthermore, in the sextuple deletion strain *rib1△/△ orf19.6263△/△ orf19.5720△/△ orf19.2751△/△ orf19.1584△/△ rut1△/△*, the cells fail to grow in riboflavin concentrations of 50 and 100 mg/L, whereas the quintuple deletion strain in which *RUT1* is present can grow, indicating its critical role in riboflavin uptake (Figure 3A and 3B). The growth trend persisted even at concentrations close to the maximum soluble amount of riboflavin (200 mg/L), confirming the importance of Rut1 in enabling growth. It appears that riboflavin can diffuse into the cell at these high concentrations, as has been observed previously (21, 28, 33). We also similarly tested the other four *ScMCH5* orthologs in quintuple deletion strains where either *ORF19.6263*, *ORF19.5720*, *ORF19.1584,* or *ORF19.2751* genes were still present but none of the strains were able to grow in lower riboflavin concentrations in the absence of *RUT1* (Supplementary Figure 3B, 3C and 3D). Again, this indicates that the four other orthologs are not involved in riboflavin uptake.

**Figure 3:**
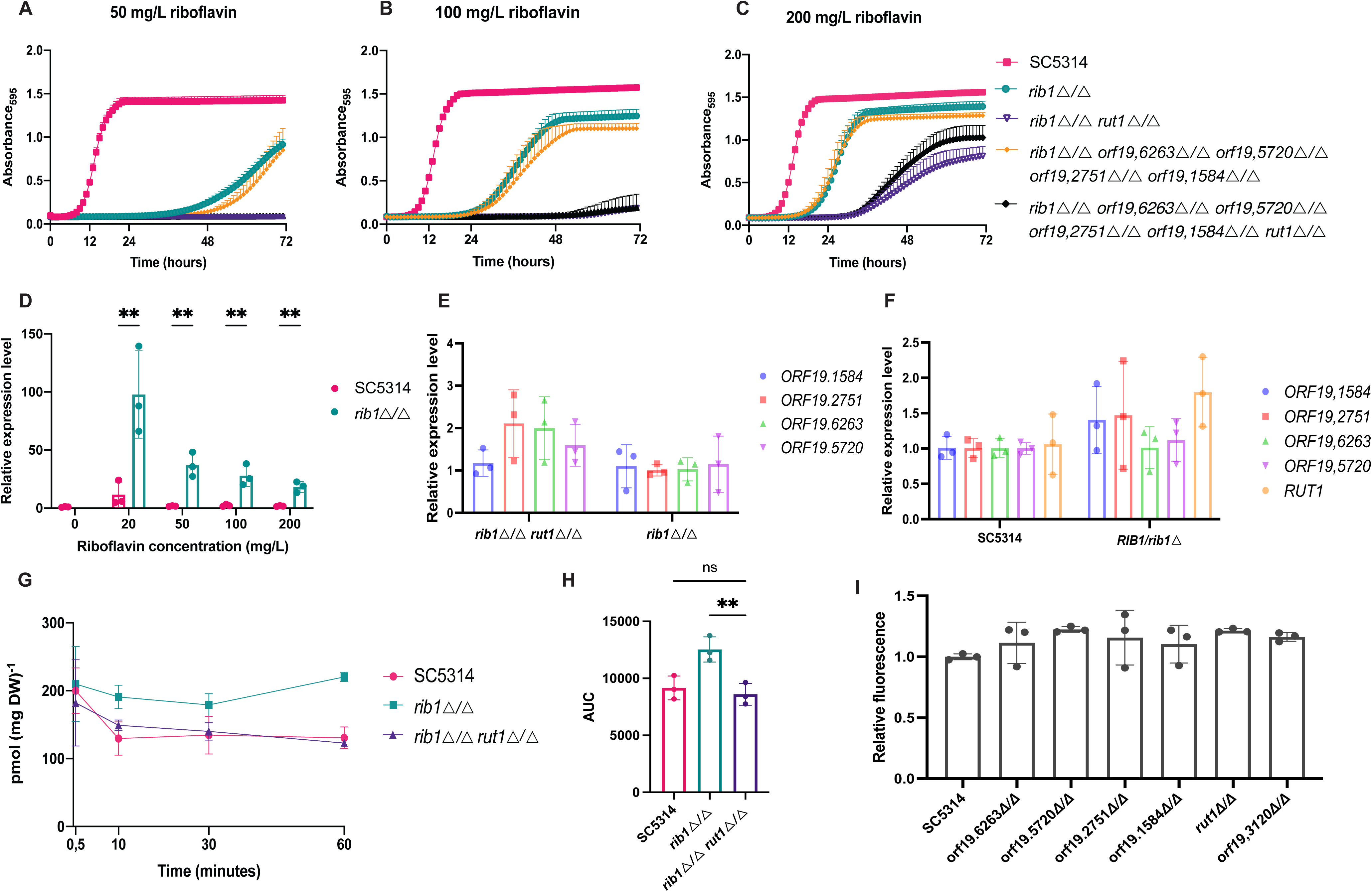
Rut1 is a major riboflavin transporter. Panels A, B and C show the growth of wild-type strain SC5314, *rib1△/△, rib1△/△ orf19.6263△/△ orf19.5720△/△ orf19.2751△/△ orf19.1584△/△* and *rib1△/△ orf19.6263△/△ orf19.5720△/△ orf19.2751△/△ orf19.1584△/△ rut1△/△* in 50 mg/L, 100 mg/L and 200 mg/L of supplemented riboflavin, respectively. (D) The relative expression level of *RUT1* in 0, 20, 50, 100, and 200 mg/L of supplemented riboflavin in the wild-type strain compared to the auxotrophic *rib1△/△* strain. There is no data for the *rib1△/△* strain as this condition is not viable. Statistical analysis was performed using 2-way ANOVA with Bonferroni’s multiple comparisons tests. **, P < 0.01. (E) The relative gene expression level of *ORF19.*1584, *ORF19.*2751, *ORF19.*6263, and *ORF19.*5720 in the auxotrophic *rib1△/△ and rib1△/△ rut1△/△* strains when grown in LoFlo medium supplemented with 100 mg/L riboflavin. (F) The relative gene expression level of *ORF19.1584, ORF19.*2751, *ORF19.*6263, *ORF19.*5720, and *RUT1* in the wild-type strain and the heterozygous *RIB1/rib1△* strain that produces reduced amounts of riboflavin. The strains were grown in LoFlo medium. (G) Riboflavin uptake in the wild-type, auxotrophic *rib1△/△, and rib1△/△ rut1△/△* strains is shown in pmol/(mg dry weight) and plotted against time. (H) Area under the curve (AUC) of three independent uptake experiments with three biological and at least two technical replicates. (I) The wild-type strain and six strains with a possible riboflavin export gene deletion were grown for 24 hours, and the fluorescence of the sterile supernatant was measured at 530 nm after excitation at 450 nm. Statistical analysis was performed by ordinary one-way ANOVA with Dunnett’s multiple comparisons test **, P < 0.01. All figures show the mean ± s.d. of three biological replicates, except for panel G, which shows the mean of three technical repeats. The biological replicates for the latter panel were performed on different days and are included in Supplementary Figure 5. We observed day-to-day variance in our radioactivity experiments as expected, but all experiments show the same trend. All gene expression data show the mean ± s.d. of three biological repeats and each biological replicate represents the mean of three technical repeats.

We investigated the relative expression level of *RUT1* at different concentrations of external riboflavin for the wild-type strain and the auxotrophic strain. Expression analysis revealed that *RUT1* is upregulated in the *rib1Δ/Δ* auxotrophic strain, especially at lower riboflavin concentrations, suggesting its induction in response to riboflavin scarcity. At 20 mg/L of external riboflavin, this putative riboflavin transporter is upregulated approximately 100-fold compared to the wild-type control strain at 0 mg/L (Figure 3D). An overview of the expression levels of all five *ScMCH5* orthologs is shown in Supplementary Figure 4. Next, we examined whether one or more of the other four *ScMCH5* orthologs would be compensatory upregulated in the absence of *RUT1,* but this was not the case, indicating that these transporters do not play a role in riboflavin uptake (Figure 3E). We then compared the expression of the five *ScMCH5* orthologs in the wild-type versus the heterozygous *RIB1/rib1△* strain, expecting that a transporter might be upregulated in the heterozygous strain since this strain produces less riboflavin, but this wasn’t the case (Figure 3F). This is most likely because the heterozygous strain still produces enough riboflavin to support growth.

To confirm that riboflavin is taken up by auxotrophic cells, we performed [^14^C] riboflavin uptake experiments. The uptake experiments consistently revealed that the *rib1△/△* strain had higher uptake than the wild-type and the *rib1△/△ rut1△/△* strain. Figure 3G shows the result of one experiment. The uptake experiments were repeated three times with three technical repeats and three biological replicates, the results of the two other experiments are shown in Supplementary Figure 5.

Figure 3H shows the area under the curve (AUC) of the uptake experiments for the three independent experiments. Despite the limited uptake efficiency for all strains, the trend was consistent across multiple experiments. In previous research by Reihl *et al.* they also observed very limited uptake for *Sc*Mch5. They were only able to observe uptake in a multi-copy plasmid *ScMCH5* overexpressing strain (28). Finally, we explored the potential involvement of the *Sc*Mch5 orthologs in riboflavin export. This was assessed by quantifying the riboflavin released into the medium (28, 34–36). We hypothesized that if a riboflavin exporter was deleted, riboflavin would no longer be secreted. However, all deletion strains were still able to secrete wild-type levels of riboflavin into the medium (Figure 3I). We also tested an additional protein, called Orf19.31200, which is orthologous to the riboflavin excretase *Cf*Rfe1 identified in *Candida famata* (37). However, it did not reveal any significant role in riboflavin secretion (Figure 3I).

According to the structure prediction by Alphafold, Rut1 is a major facilitator superfamily (MFS) domain-containing protein with twelve transmembrane domains (UniProt identifier: A0A1D8PNK9) (38–40). The predicted structure is displayed in Figure 4.

**Figure 4:**
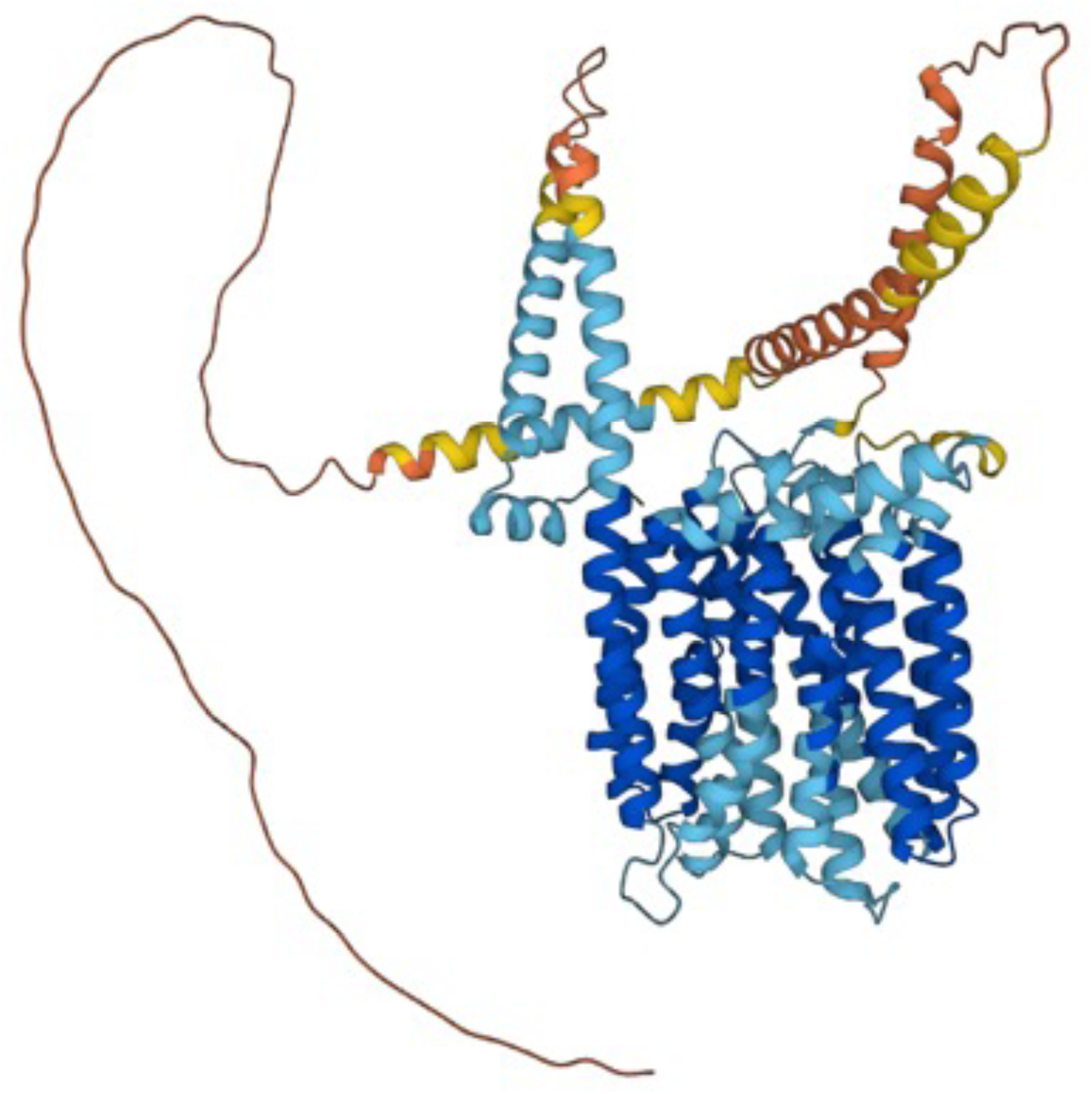
AlphaFold structure prediction of Rut1. Dark blue represents a per residue model confidence score (pLDDT) higher than 90, light blue between 70 and 90, yellow between 50 and 70, and orange represents a pLDDT score below 50.

Notably, the deletion of the putative riboflavin transporters doesn’t affect growth in LoFlo medium (Figure 5A). The same phenotype was observed at higher riboflavin concentrations (Supplementary Figure 6), indicating that *C. albicans* tends to prioritize endogenous riboflavin production over uptake. Therefore, we investigated whether *RIB1* gene expression is affected by extracellular riboflavin concentrations in a wild-type strain capable of producing riboflavin. We didn’t observe a significant decrease in gene expression but there seems to be a trend that *RIB1* is expressed less at higher riboflavin concentrations. However, we must keep in mind that these concentrations are not physiologically relevant, and the observed trend aligns with the known phenomenon of riboflavin diffusion into the cell at high concentrations. If more riboflavin enters the cell, it makes sense that there would be feedback inhibition. In conclusion, our results establish Rut1 as the major riboflavin importer in *C. albicans*, and provide additional insights into the regulation of riboflavin homeostasis in this pathogenic fungus.

**Figure 5:**
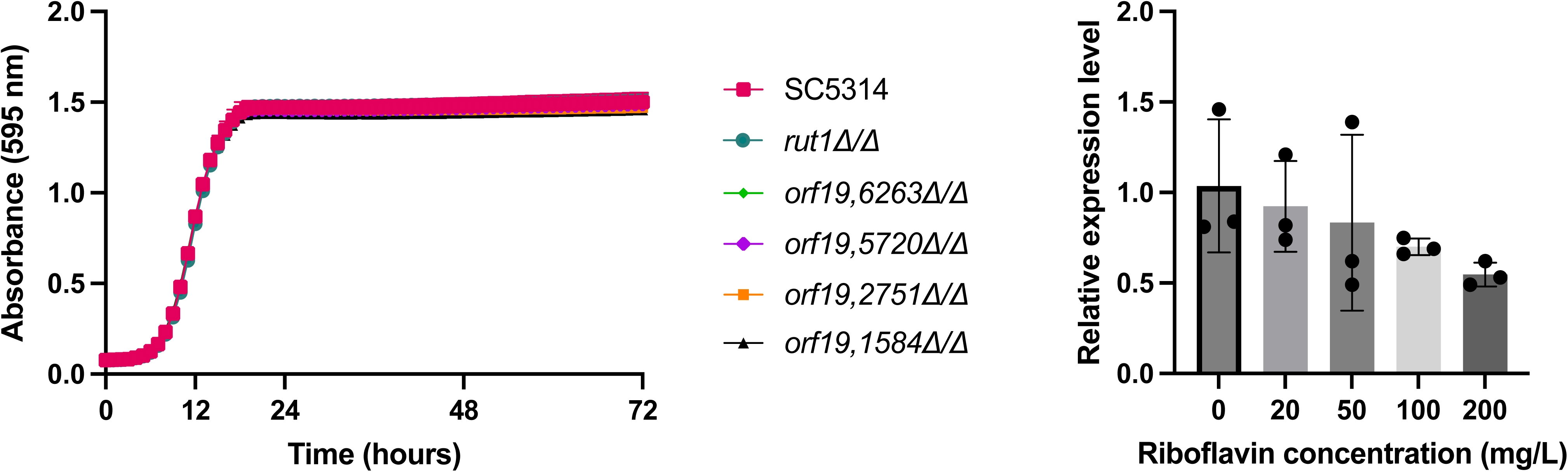
The transporters do not have a direct impact on growth. (A) The growth of different strains in LoFlo without riboflavin (B) The relative expression level of *RIB1* in the wild-type strain that was grown on different concentrations of riboflavin: 0, 20, 50, 100, and 200 mg/L. The bars show the mean ± s.d. of three biological replicates and three technical repeats. Statistical analysis was performed with an ordinary one-way ANOVA with Dunnett’s multiple comparisons test and the relative gene expression of *RIB1* is not significantly different in higher riboflavin concentration versus no riboflavin.

### Characterization of the riboflavin biosynthetic pathway in *Candida glabrata*

*Candida glabrata* is the most prevalent non-*albicans Candida* species and a common cause of candidemia. Moreover, the emergence of co-resistance to both azoles and echinocandins is of concern (41). Hence, we have also characterized the pathway in this species since it has not been previously characterized, and numerous enzymes are still uncharacterized. We searched for orthologous enzymes to the *S. cerevisiae* Rib enzymes, and identified clear orthologs for all *Cg*Rib enzymes, except for *Cg*Rib2.

We found two proteins orthologous to *Sc*Rib2: CAGL0C05291g and CAGL0K01485g, both with E-values of 0.0, indicating an exact match. The latter enzyme, known as *Sc*Pus9 in *S. cerevisiae*, resulted from the whole-genome duplication that occurred in the common ancestor of both species (42). However, it is unlikely that this gene is involved in riboflavin biosynthesis as the protein was found to be localized in the mitochondria in *S. cerevisiae,* whereas riboflavin synthesis occurs in the cytoplasm (43). In addition, the *Sc*Pus9 ortholog lacks the three conserved deaminase motifs required for the deamination activity in the riboflavin biosynthetic pathway (44). Therefore, we investigated the role of the other gene, CAGL0C05291g, in riboflavin biosynthesis.

To characterize the *CgRIB* genes in *C. glabrata*, we constructed deletion strains of the ORFs listed in Table 1. Previous research has shown that *Cg*Rib7 and *Cg*Rib1 orthologs catalyze the same reaction as their respective orthologs in *S. cerevisiae* (45, 46). Hereafter, we will refer to putative *C. glabrata* orthologs as *Cg*Rib instead of their systematic names, which are listed in Table 1.

**Table 1:** Results of protein BLAST of the *S. cerevisiae* Rib proteins against *C. glabrata*.

| S.<br>*cerevisiae* | *C. glabrata* |  |  |  |  |
| --- | --- | --- | --- | --- | --- |
|  | ID | E-value | %identity | %similarity | Associated protein |
| *ScRib1* | CAGL0F04279g/Rib1 | 4e<sup>-177</sup> | 79.6 | 88.4 | *CgRib1* |
| *ScRib2* | CAGL0C05291g | 0.0 | 65.7 | 81.2 | *CgRib2* |
|  | CAGL0K01485g | 0.0 | 60.0 | 75.6 | / |
| ScRib3 | CAGL0I07557g | $2e^{-111}$ | 72.5 | 82.6 | CgRib3 |
| ScRib4 | CAGL0B01419g/Rib4 | $2e^{-105}$ | 82.7 | 93.5 | CgRib4 |
| ScRib5 | CAGL0D05302g | $1e^{-126}$ | 72.6 | 86.5 | CgRib5 |
| ScRib7 | CAGL0F05203g/Rib7 | $2e^{-97}$ | 57.2 | 74.9 | CgRib7 |

All *CgRIB* genes were deleted in the ATCC2001 (= CBS 138) background. We monitored the growth by measuring the absorbance over a period of 72 hours. None of the *Cgrib*Δ strains were able to grow when no external riboflavin was added in the medium (Figure 6A), although candidagenome.org states that a *CgRIB2* null mutant is viable. When riboflavin was added to the medium, the growth of the *Cgrib*Δ strains was rescued. However, they did not reach the same growth as the wild-type strain when 200 mg/L of riboflavin was added, as shown in Figure 6B. To determine the riboflavin content, the fluorescence of the cell cultures was measured at 530 nm after excitation at 450 nm. As expected, none of the *Cgrib*Δ strains were able to produce riboflavin, as shown in Figure 6C. Similar to *C. albicans*, we observed that the *Cgrib*Δ strains are unable to sustain growth in RPMI medium and cells start dying over the course of 72 hours (Figure 6D). Subsequently, we examined the amount of riboflavin that was required to maintain growth in this species. To this end, we cultured a *Cgrib1*Δ mutant in varying concentrations of riboflavin (Figure 6E). The mutant was able to maintain growth at lower concentrations than *C. albicans,* starting at 10 mg/L of external riboflavin. These results indicate that the riboflavin biosynthetic pathway and its associated enzymes are necessary for maintaining viability and represent potential targets for drug development.

**Figure 6:**
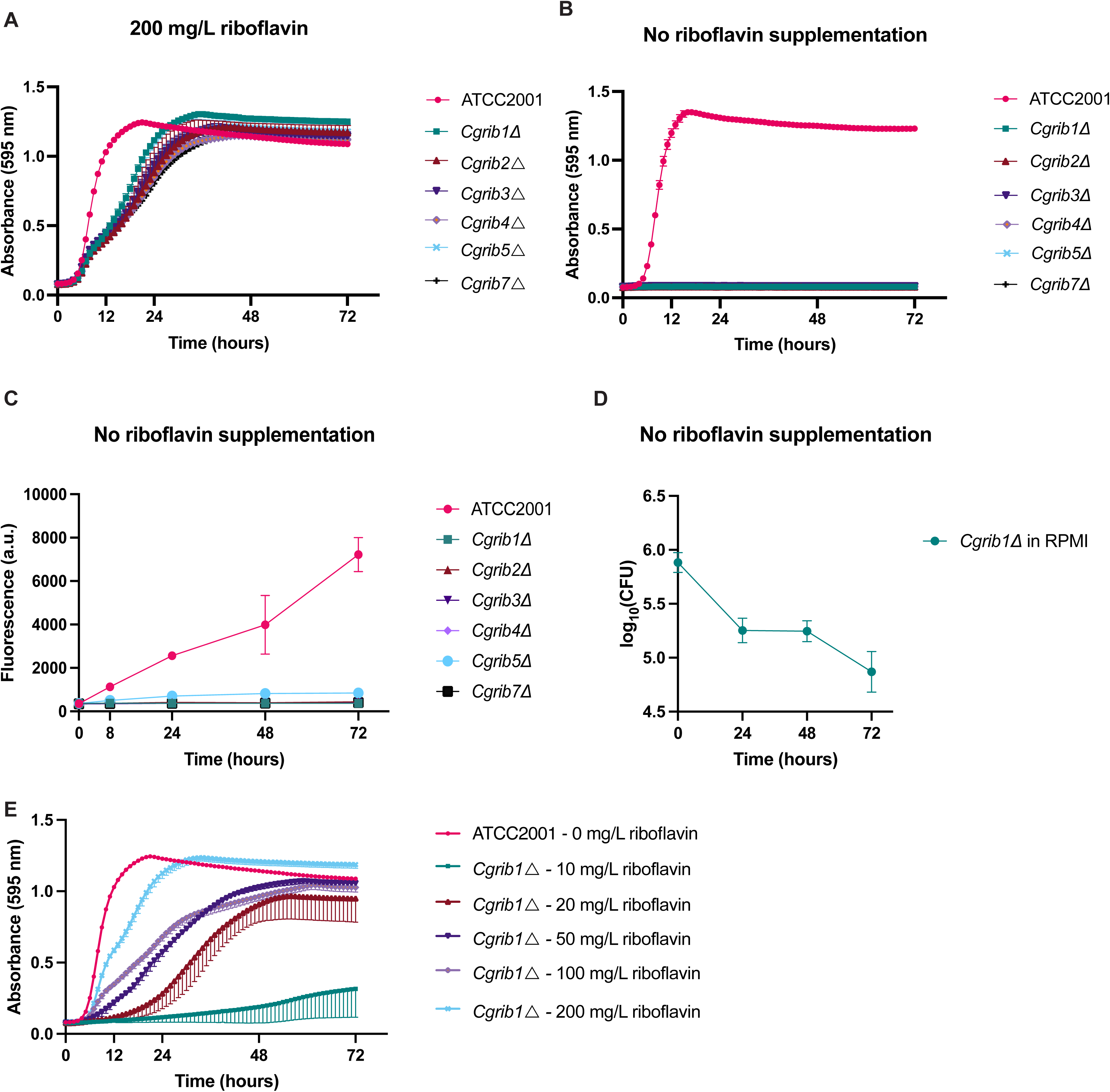
Characterization of the riboflavin biosynthetic pathway in *Candida glabrata*. (A) Deletion strains of all enzymes in the riboflavin biosynthetic pathway were cultured in a growth medium without supplemented riboflavin, and their absorbance was measured over time. (B) The strains were grown in a medium supplemented with 200 mg/L of external riboflavin (C) The strains were grown for 24h in a growth medium without external riboflavin supplemented and the riboflavin production was monitored over time by fluorescent measurements as described in the material and methods section. (D) A *RIB1* deletion strain is grown in blood-simulative RPMI medium that contains a physiological riboflavin concentration of 0,2 mg/L. CFUs were counted every 24 hours on YPD medium with 200 mg/L riboflavin to assess cell survival. (E) The growth of the *RIB1* deletion strain was monitored in LoFlo medium with different concentrations of added riboflavin. All experiments were conducted with three biological replicates, and the mean ± s.d. is presented.

### The characterization of the riboflavin biosynthetic pathway in *Saccharomyces cerevisiae*

Finally, we compared the riboflavin biosynthetic pathway of the *Candida* species with the model organism *S. cerevisiae*. Consequently, we deleted all *ScRIB* genes in the S288C background and evaluated the essentiality of each gene. Our findings revealed that all *ScRIB* genes are required for growth, as depicted in Figure 7A. This contradicts with previous reports suggesting that *Sc*Rib1 and *Sc*Rib4 were dispensable, indicating that the reaction could proceed without enzymatic catalysis (28). In contrast to *Candida* species, the growth of *ScRIB* deletion strains was fully restored at a concentration ten times lower than that of *C. albicans* and *C. glabrata*, as shown in Figure 7B. No riboflavin was detected in the cell cultures of the *ScRIB* deletion strains, as demonstrated in Figure 7C. We further examined whether the auxotrophic strains could survive in LoFlo and RPMI media and found a striking contrast. In LoFlo, the cells die in the absence of riboflavin. However, the *Scrib1*Δ mutant was able to survive in RPMI, which contains a physiological riboflavin concentration of 0.2 mg/L (Figure 7D). This suggests that when a compound inhibits the riboflavin biosynthetic pathway, *S. cerevisiae* may survive while *Candida* strains die.

**Figure 7:**
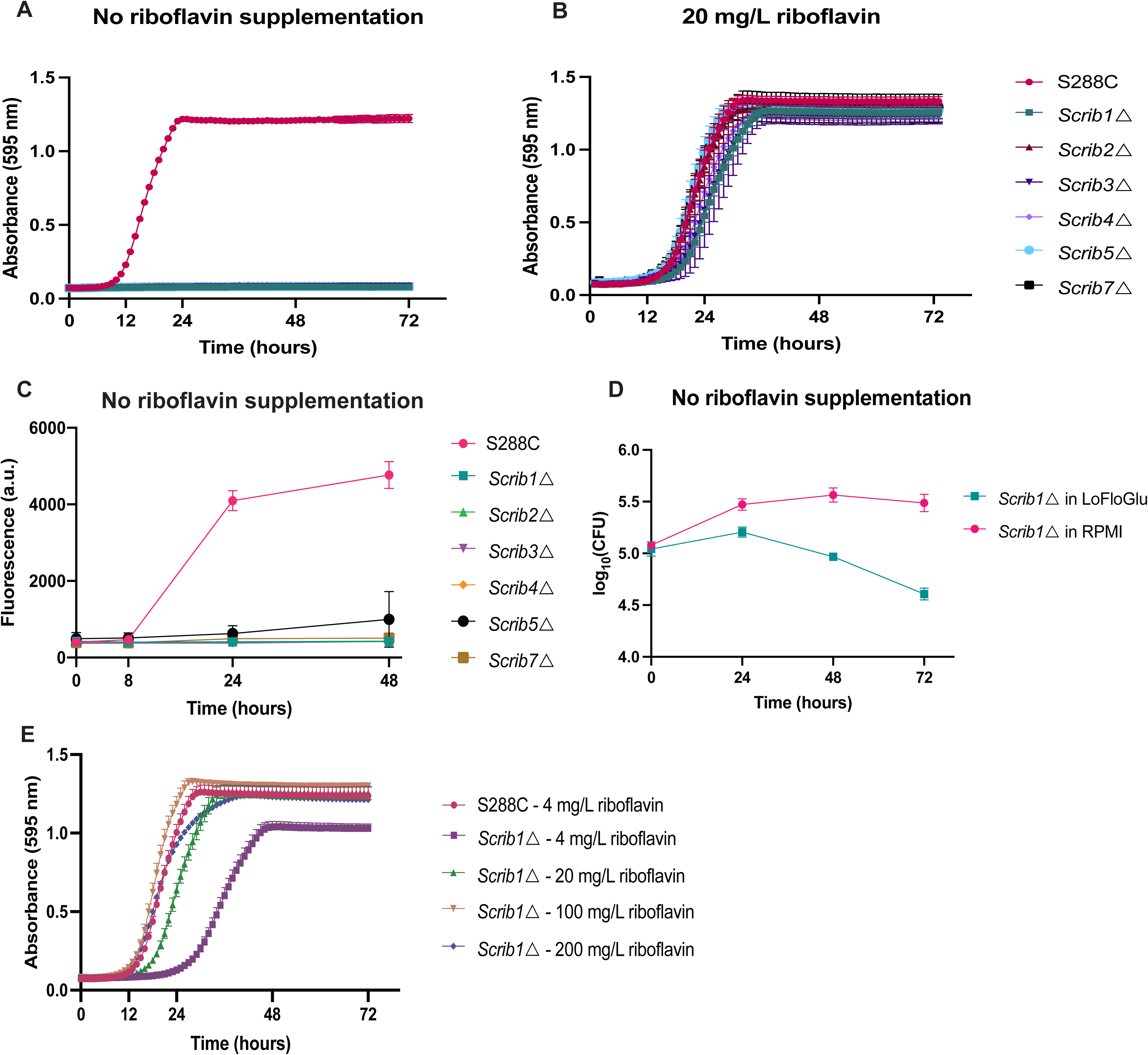
*S. cerevisiae* needs lower concentrations of riboflavin to sustain growth compared to the *Candida* species. (A) Homozygous deletion strains of all enzymes in the riboflavin biosynthetic pathway were cultured in a growth medium without supplemented riboflavin, and their absorbance was measured over time. (B) The strains were grown in a medium supplemented with 200 mg/L of riboflavin (C) The strains were grown for 24 hours in a growth medium without external riboflavin supplemented and the riboflavin production was monitored over time by fluorescent measurements. (D) A *RIB1* deletion strain was grown in LoFlo and RPMI. CFUs were counted every 24 hours on YPD medium with 200 mg/L riboflavin to assess cell survival. (E) The growth of the *RIB1* deletion strain was monitored in LoFlo medium with different concentrations of added riboflavin. All experiments were conducted with three biological replicates, and the mean ± s.d. is presented.

Finally, we examined the amount of riboflavin required for auxotrophic *S. cerevisiae* growth. We observed that this particular species can thrive at reduced riboflavin concentrations, suggesting a more efficient uptake system (Figure 8A). To examine the transport mechanism in *S. cerevisiae*, we removed *ScMCH5* in the auxotrophic *Scrib1*Δ background. Our results confirm previous reports that stated that *Sc*Mch5 is involved in riboflavin uptake, as a *Scrib1*Δ *mch5*Δ strain exhibits slower growth rates when exposed to varying riboflavin concentrations (see Figure 8B) (28).

**Figure 8:**
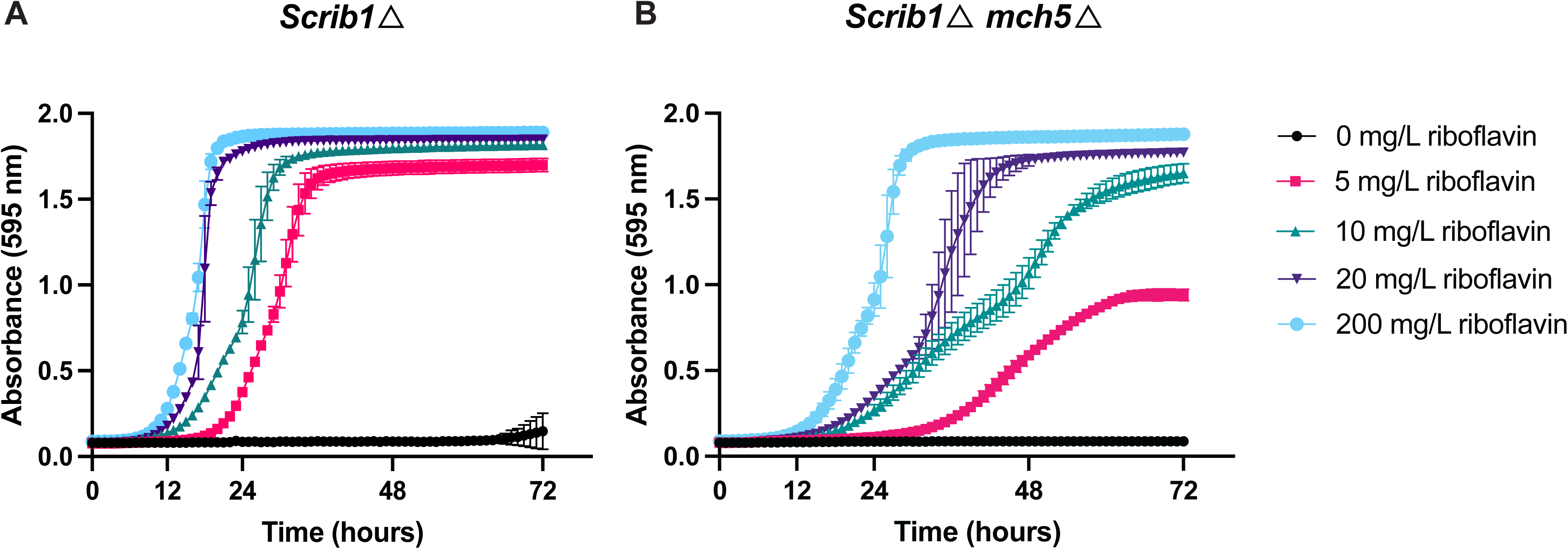
S*c*MCH5 aids in riboflavin uptake in *S. cerevisiae.* (A) The growth of *Scrib1*Δ mutant was assessed in different concentrations of supplemented riboflavin. (C) The growth of *Scrib1*Δ *mch5*Δ in different riboflavin concentrations. All experiments were conducted with three biological replicates, and the mean ± s.d. is presented.

Due to the intriguing differences in uptake between *C. albicans* and *S. cerevisiae*, we co-cultured the two *RIB1* deletion strains. Our observations revealed that the auxotrophic *S. cerevisiae* strain outperformed the *C. albicans* strain in the presence of low riboflavin concentrations of 5 mg/L, as shown in Figure 9A. This phenotype disappears at a high riboflavin concentration of 200 mg/L. When the wild-type strains are grown together in the same setup, we also do not observe this phenotype.

**Figure 9:**
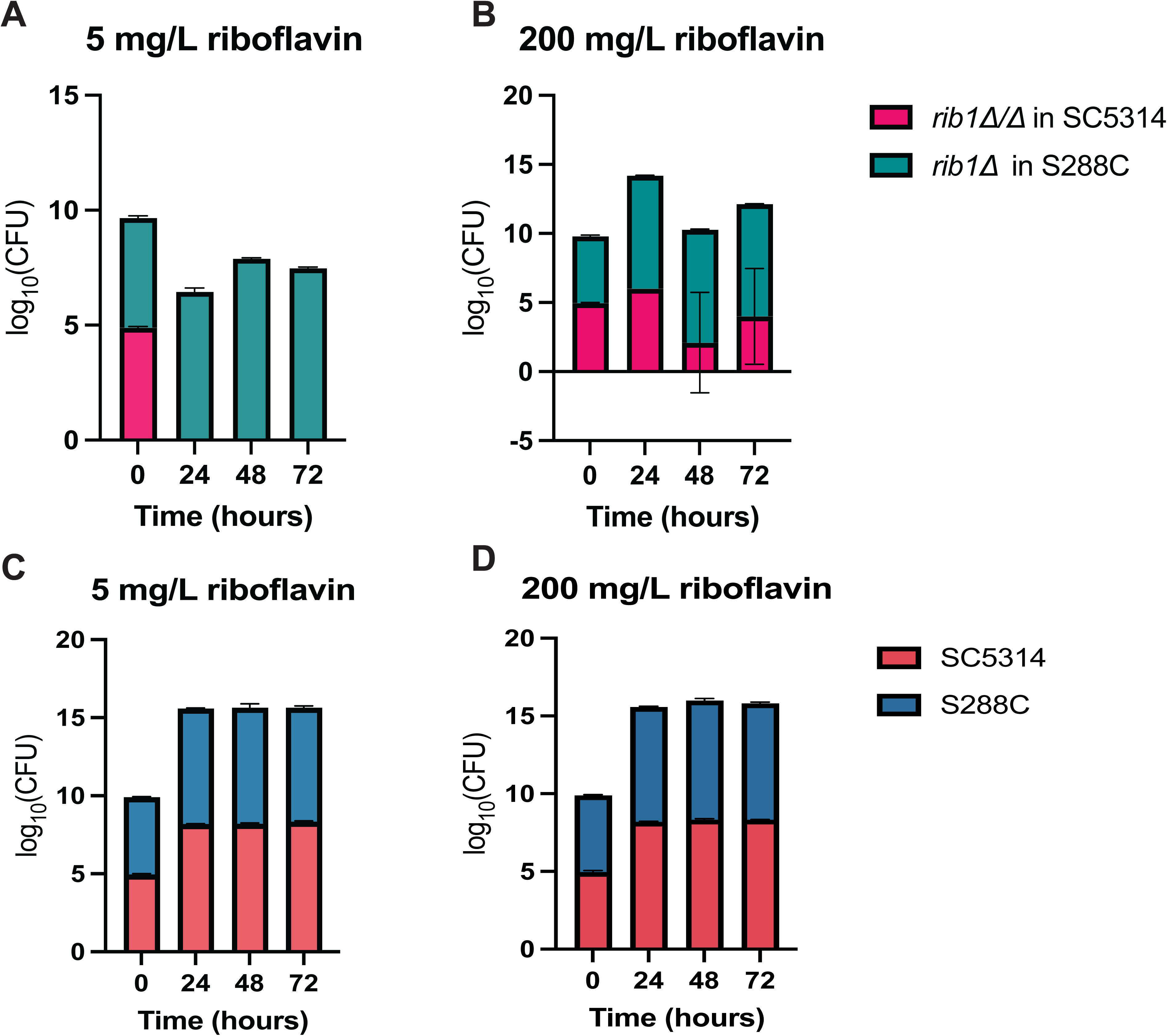
Auxotrophic *Scrib1*Δ outgrows auxotrophic *Carib1*Δ/Δ in low riboflavin concentrations. (A) Auxotrophic *RIB1* deletion strains of *C. albicans* and *S. cerevisiae* were grown together at equal starting OD_600_ in 5 mg/L of riboflavin and plated out on differentiating CHROMagar at different time points. The CFUs were counted after two days of incubation for both species and log transformed. (B) Same set-up but with 200 mg/L riboflavin. (C) The same set-up in 5 mg/L riboflavin with the wild-type strains of both species. (D) The same set-up was in 200 mg/L with the wild-type strains.

## Discussion

In this manuscript, we characterized the riboflavin biosynthetic pathway in three yeast species: *C. albicans, C. glabrata,* and *S. cerevisiae.* We demonstrate that each enzyme in the pathway is essential for growth in all three organisms. This contradicts a previous observation in *S. cerevisiae*, which reported that *Sc*Rib1 and *Sc*Rib4 were not essential, and that the reaction could occur without enzymatic catalysis (28). Although the entire pathway is essential, our investigation highlights Rib1 and Rib3 as particularly promising drug targets because the deletion of one allele reduces the amount of riboflavin produced by half and these genes are positioned at the initiation points of the two branches of the pathway leading to riboflavin synthesis. In addition, these proteins have no homology to human proteins; only one protein in the pathway, Rib2, has homology to an unidentified human RNA pseudouridylate synthase domain-containing protein (PDB: 5UBA_A). Interestingly, heterozygous deletion strains of Rib1 and Rib3 exhibited reduced riboflavin production while maintaining a growth phenotype similar to the wild-type strain. This indicates that *C. albicans* produces more riboflavin than required for growth, aligning with its classification as a flavinogenic yeast (47).

The auxotrophic strains of all species, which represent the inhibition of the pathway by a drug, were able to grow by taking up riboflavin from the environment. The growth depends on the amount of riboflavin added to the medium. Notably, there were some species-specific differences. The auxotrophic strains of *C. albicans* and *C. glabrata* require higher concentrations of around 20 and 10 mg/L of riboflavin to sustain growth, respectively, exceeding physiological levels found in the human body. To provide context, the riboflavin concentration inside human cells is typically between 0.1-0.5 mg/L and the concentration in serum or plasma ranges from 0.04 to 0.24 mg/L. The recommended daily dietary allowance of riboflavin is 1.1 to 1.3 mg, with any excess excreted in the urine (48). In contrast, *S. cerevisiae* requires a lower amount of riboflavin, suggesting a greater ability to withstand a drug targeting the pathway.

Survival experiments in blood simulative medium containing physiological riboflavin levels of 0.2 mg/L further corroborated these findings. It was observed that only auxotrophic *S. cerevisiae* could survive in this medium, while the two *Candida* species could not.

Additionally, *S. cerevisiae* was unable to survive in a medium lacking riboflavin, indicating that the uptake system in this species is more efficient than in the two *Candida* species. Our study suggests that drugs targeting the riboflavin biosynthetic pathway could be effective in inhibiting *C. albicans* and *C. glabrata* while potentially sparing *S. cerevisiae*. This has potential interesting applications as *S. cerevisiae* has probiotic properties that can inhibit *C. albicans* in certain niches (30). However, it is important to keep in mind that riboflavin concentrations in the human body can vary across tissues. Physiological riboflavin concentrations in the blood, serum, and cells are significantly lower than what is required by *C. albicans* and *C. glabrata* to sustain growth, as demonstrated by our *in vitro* experiments and confirmed *in vivo* in a systemic mouse model (29). However, the riboflavin concentration in the gut may be higher than in other tissues due to dietary intake and microbial vitamin production (17). Furthermore, our findings revealed unexpected dynamics in co-culture experiments where auxotrophic *S. cerevisiae* outcompeted auxotrophic *C. albicans* in lower riboflavin concentrations while this was not the case for higher concentrations or the wild-type strains.

Additionally, we pinpointed Orf19.4337 as the principal riboflavin uptake transporter, akin to ScMch5, a high-affinity transporter predominantly expressed under conditions of low riboflavin availability (28). We named the protein Rut1 for Riboflavin Uptake Transporter. *RUT1* is upregulated approximately 100 times more in an auxotrophic strain in 20 mg/L of riboflavin than in a wild-type strain. However, *a rib1Δ/Δ* strain can barely grow in this concentration which is 100 times higher than physiological concentrations in the human body. Due to the inefficiency of the uptake, it is unlikely that Rut1 can serve as a viable means to circumvent the mode of action of a drug that inhibits the riboflavin biosynthetic pathway. Additionally, we observed minimal uptake even in conditions where the transporter is highly upregulated. It was only slightly higher than in the wild-type strain, and the auxotrophic strain where *RUT1* is deleted. This observation was also made for *Sc*Mch5 where a multicopy plasmid was used in an auxotrophic background to measure riboflavin uptake (28). Furthermore, we found that Rut1 and the other transporters that we tested do not play a role in riboflavin export. It is possible that the export of riboflavin can occur via diffusion through the cell membrane, similar to what was observed for riboflavin import under elevated extracellular riboflavin concentrations. However, this hypothesis is tentative and necessitates further validation.

To conclude, our study provides valuable insights into the complexities of riboflavin metabolism across different yeast species, shedding light on potential drug targets and revealing intricate relationships that may impact therapeutic strategies targeting the riboflavin biosynthetic pathway.

## Material and methods

### Strains and plasmids

The *C. albicans*, *C. glabrata* and *S. cerevisiae* strains utilized in this study are listed in Supplementary Table 1. The plasmids and primers are summarized in Supplementary Tables 2 and 3, respectively.

### Cloning and transformations

Homozygous *C. albicans* strains were generated using the Hernday CRISPR system as described by Nguyen et al (49). The target-specific gRNA cassettes were amplified from pADH110 and pADH147 using the HIS-FLP split marker system (49). These were then co-transformed with the linearized *CAS9*-containing plasmid pADH99. Duplexed donor DNA, which included the upstream and downstream sequences of the target gene, was added to repair the double-stranded break induced by Cas9.

The strains were then plated on selective nourseothricin-containing plates supplemented with 200 mg/L of riboflavin. Diagnostic PCR was used to verify the colonies. The correct mutants were cultured in YP_maltose_ to induce the FLP recombinase, which removes the NAT1 marker. This allows for the reuse of the same marker and method to delete multiple genes in the same strain. The strains in which a *RIB* gene was deleted were consistently recovered and grown on YPD supplemented with 200 mg/L riboflavin. Diagnostic PCR was performed on purified DNA using primer pairs that bind to sequences within the deleted gene, which should not result in amplification, and primer pairs that bind to the region flanking the deleted gene, which should result in a short band in gel electrophoresis.

To obtain *C. albicans* reintegrant strains, the target gene was cloned into the integrative CIp10-NAT1 plasmid using NheI and AatII restriction enzymes (50, 51). The constructed plasmids were linearized with StuI, and transformed into the corresponding deletion strain using the lithium acetate method, as described by Gietz et al. (52). The correct integration was verified by diagnostic PCR, and the copy number was determined using qPCR. Transformants with two integrations of the target gene were selected.

Heterozygous *C. albicans* strains were created using the *SAT1* flipper tool, as described by Reuß et al. (31). A plasmid housing a deletion cassette with 500 bp homologous regions on both sides of the *RIB* genes was engineered. These homologous regions were amplified from the upstream and downstream regions of the *RIB* gene using genomic DNA from the wild-type strain. The backbone utilized was pSFS2n, with ApaI and XhoI restriction enzymes applied for the introduction of the upstream fragment, and NotI and SacI employed to introduce the downstream homologous fragment regions.

After extracting the correct Gibson (NEBuilder^®^ HiFi DNA Assembly) ligated plasmid from competent *E. coli* cells, we cut the plasmid using ApaI and SacI and transformed it into *C. albicans* wild-type cells with the lithium acetate method (52). The transformed cells were then plated on YPD plates containing 200 μg/mL nourseothricin (WERNER BIO). Subsequently, the cassette is removed from the genome through FLP-mediated excision.

*C. glabrata* deletion strains were constructed in the ATCC2001 (CBS138) background by replacing the GOI through homologous recombination with a deletion cassette (53). The cassette contains a *NAT1* flanked by FRT sites and 100 bp of the up-and-downstream regions flanking the GOI. These cassettes were amplified from pYC44 using PCR and then linearized with EciI. The transformed cells were plated on selective plates containing nourseothricin and supplemented with 200 mg/L of riboflavin. The colonies were verified with diagnostic PCR. Next, the correct strains were transformed with pLS10, a plasmid that contains a flippase enzyme that removes the nourseothricin cassette. The strains were plated on selective plates containing hygromycin and supplemented with 200 mg/L of riboflavin. The correct removal was verified using PCR. The pLS10 plasmid was removed by growing the strains overnight on a non-selective YPD riboflavin medium, and the removal was confirmed by restreaking the colonies on YPD with and without hygromycin.

To delete the *RIB* genes in the laboratory haploid *S. cerevisiae* CEN.PK 113-7D (S288C) strain, the KanMX cassette (a G418 resistance gene) from the low copy plasmid pTOPO-A1-G2-B-KanMX-P-G2-A2 was amplified using primers with 69 bp homologous tails, aligning with the specific GOI regions. Subsequent transformations of the cassettes were carried out using an adapted version of the LiAc/SS-DNA/PEG system described by Gietz et al (54). To create a double gene deletion, we used a pTOPO-A1-G2-B-KanMX-P-G2-A2 plasmid and selected the correct transformants on geneticin.

We obtained at least three biological replicates for every strain and stored all verified deletion strains at −80°C in stock medium.

### Growth conditions: media and chemicals

The strains were cultivated in LoFlo medium, which contains 2% glucose and no riboflavin. It has a very low fluorescence background, making it ideal for fluorescent readouts. This medium consists of 0,19% yeast nitrogen base without amino acids, folic acid, and riboflavin (Formedium), 0,079% complete supplement mixture (MP Biomedicals), and 2% glucose, which was added after autoclaving. The pH was adjusted to 5.5. In some experiments, glucose was replaced by 2% glycerol. RPMI (1,04% RPMI 164 (Thermo Fisher Scientific), 3,453% morpholine propane sulfonic acid (MOPS) at pH 7) and YPD (1% yeast extract, 2% bacteriological peptone, and 2% glucose) were used when mentioned in the text. Riboflavin at a concentration of 200 mg/L was always supplemented to the medium to make precultures and was added to experiments when noted in the text. *C. albicans* and *S. cerevisiae* were incubated at 30°C whereas *C. glabrata* was grown at 37°C.

### Copy number determination by qPCR

The copy number of re-integrant strains was determined by extracting the genomic DNA and performing quantitative PCR (qPCR) using GoTaq polymerase (Promega) and a StepOnePlus real-time PCR machine (ThermoFisher). The PCR reaction contained 2.5 ng of genomic DNA. The CaACT1 promoter region present in the CIp10 plasmid was amplified along with three reference genes: Ca18S, CaACT1, and CaTEF1. The primers are listed in Supplementary Table 3. The data were analyzed using qBasePlus software (Biogazelle).

### Growth analysis

Growth was assessed by measuring absorbance at 595 nm over time in a Multiskan FC microplate photometer (Thermo Scientific) using flat-bottom 96-well plates (Bioké) and intermittent (10-minute interval) pulsed shaking (medium strength, 1 minute). The cells were washed three times in phosphate-buffered saline (PBS), and growth was subsequently measured at 30°C with a starting OD of 0.01. The growth curves represent the average of three biological replicates for each strain with ± s.d..

### Riboflavin quantification by fluorescence measurement

Prior to measurement, cells were washed three times with PBS and the OD_600_ was adjusted to 0.01 in LoFlo growth medium. The resulting solution was added to a black-walled flat-bottom 96-well plate (Greiner) and incubated at 30°C in a shaking plate incubator protected from the light. Riboflavin was quantified through fluorescence measurements in the Synergy H1 Hybrid multimode microplate reader (BioTek) using emission at 530 nm upon excitation at 450 nm. The growth of the same cultures could be measured in the same machine at an absorbance of 600 nm.

### Intracellular riboflavin measurements using Flow cytometry

Overnight precultures of *C. albicans* in LoFLo medium supplemented with 200 mg/L of riboflavin underwent three washing steps and were subsequently loaded into a U-shaped 96-well plate (Greiner) at an OD_600_ of 0.05. Intracellular fluorescence, specifically green fluorescence (GRN B-Hlog) that corresponds to the riboflavin wavelengths, was measured using the Guava easyCyte Flow Cytometer (Cytek). 10 000 cells were excited with a 488 nm laser, and the emitted light was captured through a 525/30 bandpass filter. The obtained data was analyzed using Flowjo software and was gated to exclude debris or aggregates. The graphs illustrate the median values derived from three biological repeats.

### Survival of auxotrophic *RIB* deletion strains

The survival of auxotrophic *RIB* deletion strains was assessed by first washing the overnight preculture three times in PBS. Then, 1*10^5^ cells/mL were placed in test tubes with RPMI-MOPS 1640 growth medium or LoFlo medium at 30°C for *C. albicans* and *S. cerevisiae* and 37°C for *C. glabrata*. At time point zero and every 24 hours, 100 µL of culture was diluted in growth medium to a final concentration of 1000 cells/mL. Subsequently, 100 µL of this diluted culture was plated on a YPD plate supplemented with 200 mg/L riboflavin. Two days later, CFUs were counted (approximately 100 cells per plate).

### RNA extraction and gene expression analysis using qRT-PCR

Overnight precultures were grown in 50 mL LoFLo medium at an OD_600_ of 0,1 at 30°C. For some experiments, different levels of riboflavin were also added to the medium. The cells were grown until they reached an OD_600_ of 1, then cells were harvested and washed in ice-cold water. Subsequently, the cells were resuspended in TRIzol (ThermoFisher) and broken-down using glass beads and a FastPrep machine (MP Biomedicals). Pure DNA was isolated using chloroform and isopropanol, and three washed in 70% ethanol. The RNA concentration was measured using a NanoDrop spectrophotometer (ND-1000; Isogen Life Science), and the RNA was treated with DNase (New England Biolabs). Next, the RNA was converted to complementary DNA using an iScript cDNA synthesis kit (Bio-Rad). Quantitative real-time PCR (qRT-PCR) was performed with the GoTaq polymerase (Promega) and a StepOnePlus real-time PCR device (ThermoFisher). The primers used are listed in Supplementary Table 3, along with the sequences of the three reference gene primers: *Ca18S*, *CaACT1,* and *CaTEF1.* The data was analyzed using qBasePlus software and the statistical analysis was performed with GraphPad Prism. The data was log_2_(Y) transformed and statistical analysis was conducted using ordinary one-way ANOVA with Dunnett’s multiple comparisons test or a t-test, depending on the experiment. Each experiment included three biological and three technical repeats, and the graphs display the untransformed data with mean ± s.d.

### Riboflavin uptake

Transport assays were performed similary to those described by Soares-Silva et al. (55) but optimized to measure riboflavin uptake. Wild-type, *Carib1Δ/Δ,* and *Carib1Δ/Δ rut1Δ/Δ* strains were grown overnight in 3 mL LoFlo medium with 200 mg/L of riboflavin at 30°C. The OD_600_ was adjusted to OD 2 in 50 mL LoFlo medium with 200 mg/L of riboflavin, and the cells were grown overnight to obtain sufficient cells. The following day, the cells were washed three times with PBS. The _OD600_ was adjusted to 0.5 and the cells were deprived of riboflavin for 4 hours by growing them in LoFLo medium supplemented with 10 mg/L riboflavin. Cells were then harvested on ice, washed twice, and resuspended in 1 mL of LoFlo medium. The dry weight was determined by transferring two times 100 µL of this culture to a filter paper, which was dried for two days at 50°C (dry weight should be between 20 and 40 mg/mL). Reaction mixtures were prepared by adding 30 µL of cells to an Eppendorf tube together with 60 µL of LoFlo medium. The cells were incubated for 2 minutes at 30°C, and 10 µL of 0,1 mM [^14^C] riboflavin (Campro Scientific; 0,37MBq with an activity of 30 mCi/mmol) was added to start the reaction, and the mixture was vortexed briefly. After 30 seconds, 10 minutes, 30 minutes, and 1 hour, 1 mL of ice-cold Milli-Q water was added to the cells to stop the reaction. The cells were always kept on ice and were collected by centrifugation at 7000 rpm for 2 minutes at 4°C. Then, 1 mL of the supernatant was discarded before adding of 1 mL ice-cold Milli-Q water. The cells were centrifuged again, and 1 mL of the supernatant was removed. This last step was repeated to ensure that all riboflavin was removed from the samples. Finally, all supernatant was removed from the samples and 1 mL of scintillation liquid (Lumagel Safe; PerkinElmer) was added. The mixture was vortexed and resuspended until the pellet was completely dissolved. Riboflavin uptake was measured by placing the Eppendorf tubes in scintillation vials in the scintillation counter (Hidex). Each experiment was repeated three times with three technical replicates and a different biological repeat of the mutant strains each time. The specific activity of the [^14^C] riboflavin solutions was measured in every experiment by adding 10 µL of [^14^C] riboflavin to an Eppendorf tube with 1 mL of scintillation fluid.

## Author contributions

Conceptualization: J.N.; Data curation: J.N.; Formal analysis: J.N.; Funding Acquisition: P.V.D..; Investigation: J.N., D.V., and A.P. Methodology: J.N., L.D., P.V.D. Resources: P.V.D.; Supervision: J.N., L.D., and P.V.D.; Visualization: J.N.; Writing – original draft: J.N.; Writing – review & editing: J.N., A.P., L.D., and P.V.D.

## Acknowledgments

The authors thank Rudy Vergauwen for his valuable insights and feedback. J.N. and A.P. were supported by personal grants from the Fund for Scientific Research Flanders (FWO grant 1S18121N and 1S19219N). This work was supported by the KU Leuven Research Council (grant # C14/22/075) to P.V.D.

## Declaration of interest

The authors declare no conflict of interest.

